# NVX-CoV2373 vaccine protects cynomolgus macaque upper and lower airways against SARS-CoV-2 challenge

**DOI:** 10.1101/2020.08.18.256578

**Authors:** Mimi Guebre-Xabier, Nita Patel, Jing-Hui Tian, Bin Zhou, Sonia Maciejewski, Kristal Lam, Alyse D. Portnoff, Michael J. Massare, Matthew B. Frieman, Pedro A. Piedra, Larry Ellingsworth, Gregory Glenn, Gale Smith

## Abstract

There is an urgent need for a safe and protective vaccine to control the global spread of SARS-CoV-2 and prevent COVID-19. Here, we report the immunogenicity and protective efficacy of a SARS-CoV-2 subunit vaccine (NVX-CoV2373) produced from the full-length SARS-CoV-2 spike (S) glycoprotein stabilized in the prefusion conformation. Cynomolgus macaques (*Macaca fascicularis*) immunized with NVX-CoV2373 and the saponin-based Matrix-M adjuvant induced anti-S antibody that was neutralizing and blocked binding to the human angiotensin-converting enzyme 2 (hACE2) receptor. Following intranasal and intratracheal challenge with SARS-CoV-2, immunized macaques were protected against upper and lower infection and pulmonary disease. These results support ongoing phase 1/2 clinical studies of the safety and immunogenicity of NVX-CoV2327 vaccine (NCT04368988).

**Highlights:**

- Full-length SARS-CoV-2 prefusion spike with Matrix-M1™ (NVX-CoV2373) vaccine.
- Induced hACE2 receptor blocking and neutralizing antibodies in macaques.
- Vaccine protected against SARS-CoV-2 replication in the nose and lungs.
- Absence of pulmonary pathology in NVX-CoV2373 vaccinated macaques.

## 1. INTRODUCTION

There is an urgent need for a safe and effective severe acute respiratory syndrome coronavirus 2 (SARS-CoV-2) vaccine to prevent coronavirus disease 2019 (Covid-19). We have developed a recombinant nanoparticle vaccine constructed from the full-length, wild-type SARS-CoV-2 spike glycoprotein (GenBank gene sequence MN908947, nucleotides 21563-25384) optimized for the baculovirus-*Spodoptera frugiperda* (Sf9) insect cell expression system [1]. In mice and nonhuman primates (NHP), NVX-CoV2373 with a Matrix-M1 saponin-based adjuvant induced high titer anti-spike IgG that blocks binding to the hACE2 receptor, neutralize wild type virus, and protects mice against SARS-CoV-2 challenge with no evidence of vaccine-associated enhanced respiratory disease. NVX-CoV2373 vaccine also induces polyfunctional CD4^+^ T-cell responses of IFN-γ, IL-2, and TNF-α biased towards a Th1 phenotype, and generates antigen-specific germinal center B cells in the spleen. Safety and immunogenicity NVX-CoV2327 vaccine is currently under evaluation in humans (NCT04368988) and primary safety and immunogenicity outcomes described [2]. We evaluate in the current study NVX-CoV2373 vaccine immunogenicity, induction of receptor blocking, and neutralizing antibodies compared to levels in human COVID-19 convalescent sera. And in a nonhuman primate challenge model, protection against upper and lower virus replication and pulmonary disease.

## 2. MATERIALS AND METHODS

### 2.1. Cell lines, virus, antibody reagents, and receptors

Vero E6 cells (ATCC, CRL-1586) were maintained in Minimal Eagles Medium (MEM) supplemented with 10% fetal bovine serum, 1% glutamine and 1% penicillin and streptomycin. The SARS-CoV-2 (WA-1, 2020) isolated was obtained from the Center for Disease Control and stock virus prepared in Vero E6 cells. Histidine-tagged hACE2 receptor was purchased from Sino Biologics (Beijing, CN). Rabbit anti-SARS-CoV spike protein was purchased form Biodefense and Emerging Infections Research Resources Repository (BEI Resources, Manassas, VA).

### 2.2. Recombinant SARS-CoV-2 spike protein

NVX-CoV2327 was codon optimized synthetically produced from the full-length S glycoprotein gene sequence (GenBank MN908947 nucleotides 21563-25384) for expression in *Spodoptera frugiperda* (Sf9) cells (GenScript Piscataway, NJ, USA) as describe [1]. Briefly, the S1/S2 furin cleavage site 682-RRAR-685 was modified 682-QQAQ-685 and two proline substitutions introduced at positions K986P and V987P (2P) to stabilize the full-length SARS-CoV-2 S [3].

### 2.3. Animal ethics

The in-life portion of the study was conducted at BIOQUAL, Inc (Rockville, MD). Female and male, > 3 years old at study initiation, cynomolgus macaques (*Macaca fascicularis*) were obtained from Primgen, Inc (Hines, IL) and maintained at BIOQUAL, Inc for the entire in-life portion of the study. BIOQUAL, Inc. is accredited by AAALACC International. Animals were maintained and treated according to the Institutional Biosafety Committee guidelines and the study was pre-approved by the Institutional Animal Care and Use Committee (IACUC). The study was conducted in accordance with the National Institutes of Health Guide for Care and Use of Laboratory Animals (NIH publication 8023, Revised 1978).

### 2.4. Cynomolgus macaque immunization

Cynomolgus macaques >3 years old (n=4/group) at study initiation received 5 or 25 μg NVX-CoV2327 with 50 μg Matrix-M1 (Novavax AB, Uppsala, Sweden) administered in 500 μL in the thigh muscle in two doses spaced 21 days apart. A separate group was immunized with a factional dose (2.5 μg) NVX-CoV2373 with 25 μg Matrix-M1 in two doses spaced 21 days apart and a placebo group received formulation buffer. Serum was collected before immunization on Day 0, Day 21 just prior to the second immunization, and Day 33.

#### Anti-spike (S) IgG ELISA

Anti-SARS-CoV-2 spike (S) protein IgG ELISA titers were measured as described [1]. Anti-S IgG EC_50_ titers were calculated by 4-parameter fitting using SoftMax Pro 6.5.1 GxP software. Individual animal anti-S IgG EC_50_ titers, group geometric mean titers (GMT) were plotted using GraphPad Prism 7.05 software.

### 2.5. Inhibition of hACE2 receptor binding and neutralization

Antibodies that block binding of hACE2 receptor to the S-protein and neutralize in a cytopathic effect assay (CPE) in Vero E6 cells were measured as described previously as the serum titer that blocks 100% CPE [1]. Serum antibody titer at 50% binding inhibition (IC_50_) of hACE2 to SARS-CoV-2 S protein was determined in the SoftMax Pro program. Individual animal hACE2 receptor inhibiting titers, mean titers, and SEM were plotted using GraphPad Prism 7.05 software. Neutralizing antibody titers were determined as the dilution of serum that inhibited 100% of CPE (CPE_100_) at 3 days post infection of Vero E6 cells in a 96 well plate format.

### 2.6. SARS-CoV-2 challenge procedure

The virus challenge study was done at BIOQUAL, Inc. within a BSL-3 containment facility. SARS-CoV-2 generated from isolate 2019-nCoV/USA-WA1/2020 was received from BEI Resources (NR-52281; lot # 70033175) and expanded in Vero E6 cells for challenge stock generation. Animals were sedated and challenged with a targeted total dose of 1.04 × 10^4^ pfu SARS-CoV-2 by intranasal (IN) and intratracheal (IT) in a volume of 0.25 mL each route. BAL and nasal swabs were collected 2- and 4-days post challenge. Necropsy was performed 7 days following challenge and lung tissues collected for histopathology.

### 2.7. RNA subgenomic RT-PCR

The subgenomic viral mRNA (sgRNA) was measured in macaque bronchoalveolar lavage (BAL) and nasal swabs collected 2- and 4-days post challenge using RT-PCR as described [4]. To generate a standard curve, the SARS-CoV-2 E gene sgRNA was cloned into a pcDNA3.1 expression plasmid. The insert was transcribed using an AmpliCap-Max T7 High Yield Message Maker Kit (Cellscript, Madison, WI) to obtain RNA for standards. Prior to RT-PCR, samples collected from challenged animals or standards were reverse-transcribed using Superscript III VILO (Invitrogen) according to the manufacturer’s instructions. A Taqman custom gene expression assay (ThermoFisher Scientific, Rockville, MD) was designed using the sequences targeting the E gene sgRNA. Reactions were carried out on a Quant Studio 6 and 7 Flex Real-Time PCR System (Applied Biosystems, Foster City, CA) according to the manufacturer’s specifications. Standard curves were used to calculate sgRNA in copies per mL. The quantitative assay was sensitive to 50 copies per mL.

### 2.8. Human COVID-19 convalescent serum

Convalescent serum samples (n=32) were provided by Dr. Pedro A Piedra (Baylor College of Medicine, Houston, TX, USA). Samples were collected from COVID-19 individuals 18-79 years of age 4-6 weeks after testing positive for SARS CoV-2. Symptoms ranged from asymptomatic, mild to moderate symptoms, to severe symptoms requiring hospitalization. Sera were analyzed for anti-SARS-CoV-2 S IgG, hACE2 receptor inhibition, and virus neutralizing antibody titers.

### 2.9. Histopathology

Animals were euthanized 7-days following SARS-CoV-2 challenge (Day 42) and lung tissues collected. Tissue were prepared for histologic examination by Experimental Pathology Laboratories, Inc. (EPL, Sterling, VA). The lungs were fixed with 10% formalin, paraffin embedded, and sections stained with hematoxylin and eosin (H&E) for histological examination. Slides were examined for total inflammation, periarteriolar, and peribronchiolar inflammation and epithelial cell denuding.

## 3. RESULTS

### 3.1. Immunogenicity of NVX-CoV2373 in nonhuman primates compared to COVID-19 convalescent human sera

Macaques immunized with the prime/boost regimen of 2.5, 5, and 25 μg NVX-CoV2373 with 25 μg in the low and 50 μg Matrix-M1 adjuvant in the two higher doses induced anti-S IgG (EC_50_) antibodies at Day 21 after a single dose (GMT = 7,810, 22,386 and 21,472, respectively). Two weeks following a booster immunization, anti-S IgG EC_50_ titers increased to GMT EC_50_ = 163,036, 335,017 and 469,739, respectively (Figure 1A). In contrast, SARS-CoV-2 anti-S antibody in convalescent human sera was 6.9- to 14.2-fold less with at GMT EC_50_ of 23,614 (Figure 1B). And, hACE2 receptor inhibition titers of 649, 1,410, and 1,320 in 2.5, 5, and 25 µg NVX-CoV2373 dose groups respectively were 5.2 – 11.2-fold higher than in convalescent sera (Figure 1C). Finally, SARS-CoV-2 GMT neutralization antibody titers of 17,920 - 23,040 CPE_100_ in immunized macaques, were 7.9 – 10.1-fold higher than in convalescent sera (Figure 1D).

**Figure 1.**
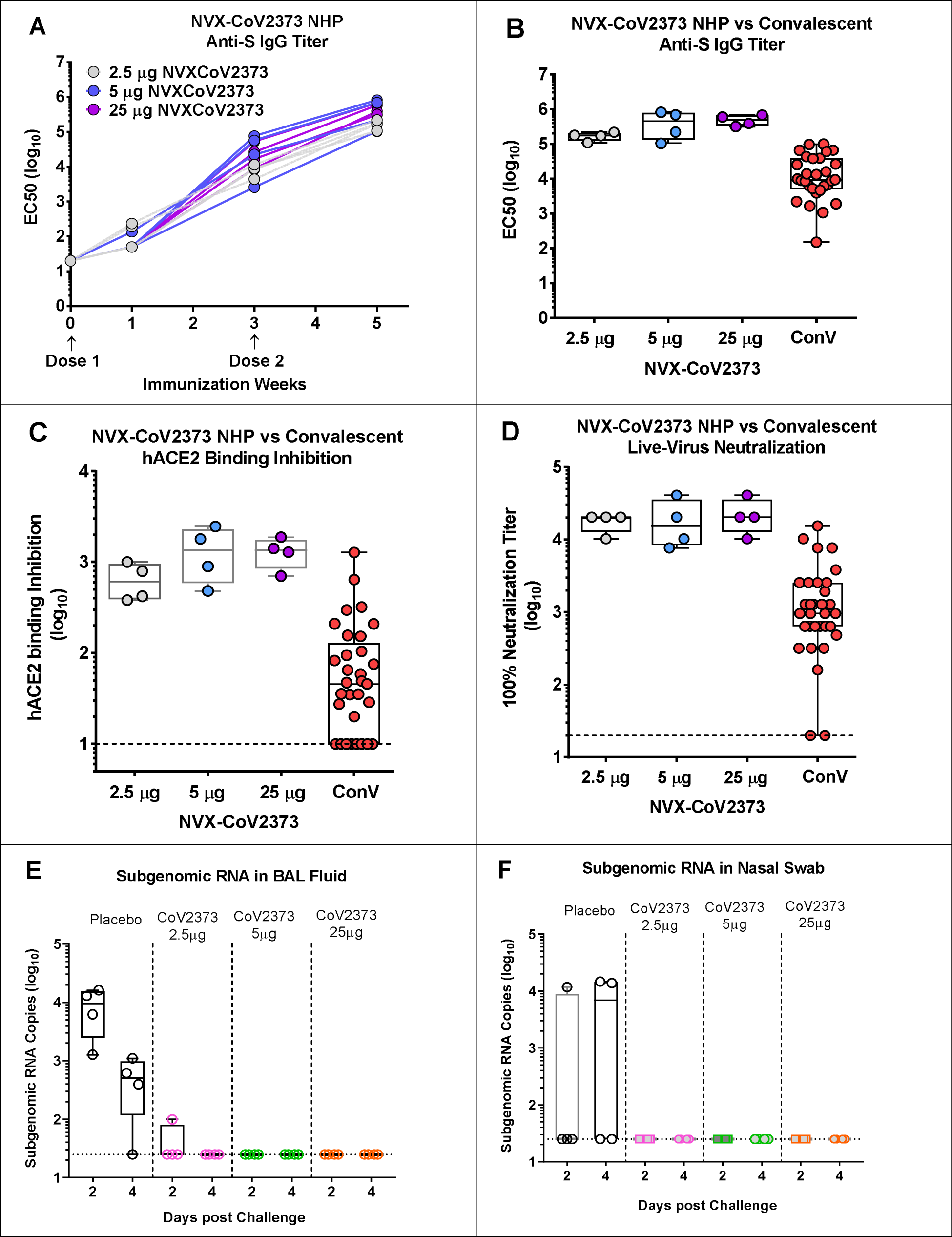
Immunogenicity of NVX-CoV2373 vaccine in cynomolgus macaques. (**A**) Groups of cynomolgus macaques (n = 4 per arm) were immunized weeks 0 and 3 with 2.5 μg NVX-CoV2373 with 25 μg Matrix-M1 or 5 μg or 25 μg NVX-CoV2373 with 50 μg Matrix-M1. Anti-spike EC_50_ IgG titers were measured weeks 0, 1, 3, and 5. Lines indicate anti-spike IgG titers for individual macaques in each group. (**B**) Anti-spike EC_50_ IgG serum titers week 5 in NVX-CoV2373 immunized NHP compared to anti-S EC_50_ IgG titers in convalescent human sera. (**C**) ACE2 inhibition IC_50_ serum titers week 5 NVX-CoV2373 immunized macaques compared to ACE2 inhibition titers in convalescent human sera, (**D**) Neutralization CPE_100_ titers against wild type SARS-CoV-2 virus week 5 NVX-CoV2373 immunized macaques compared to neutralization CPE_100_ titers in convalescent human sera, (**E**) Subgenomic RNA copies in BAL fluid days 2 and 4 post challenge SARS-CoV-2 virus in placebo and NVX-CoV2373 immunized macaques. (**F**) Subgenomic RNA copies in nasal swab samples days 2 and 4 post challenge with SARS-CoV-2 virus in placebo and NVX-CoV2373 immunized macques. Dashed horizontal line indicates the limit of detection (LOD). ConV: Convalescent serum. BAL: bronchoalveolar lavage.

### 3.2. Viral load in nasal swabs and BAL

To evaluate the potential efficacy of NVX-CoV2373 vaccine, macaques were challenged with SARS-CoV-2 virus in upper and lower airways. Macaques in the placebo group had 9,131 sgRNA copies/mL in the BAL at 2 days post challenge and remained elevated at day 4 except for one animal. In contrast, immunized animals had no detectable sgRNA in BAL fluid other than one animal in the low dose group at day 2 which cleared replicating virus RNA by day 4 (Figure 1E). Half of the controls had ~4 log10 of virus sgRNA copies in nasal swabs and in contrast, no detectable sgRNA was in the nose of NVX-CoV2373 vaccinated animals (Figure 1F).

### 3.3. Lung pathology

Lung tissues were collected from all animals at 7 days post challenge and sections examined for pathologic changes within the upper and lower airways. Consistent with previous reports of SARS-CoV-2 infection in rhesus macaques [5–10] placebo control animals had moderate to severe inflammation that involved the mucosa of the bronchi, perivascular mononuclear infiltrate with mixed infiltrates of macrophages and neutrophils within the alveoli. In contrast, there was little, or no inflammation observed in the lungs of macaques immunized with NVX-CoV2373 vaccine 7 days post challenge (Figure 2). These findings were consistent the absence of sgRNA in BAL fluids and nasal swabs of vaccinated animals by day 4 post challenge (Figure 1E and 1F).

**Figure 2.**
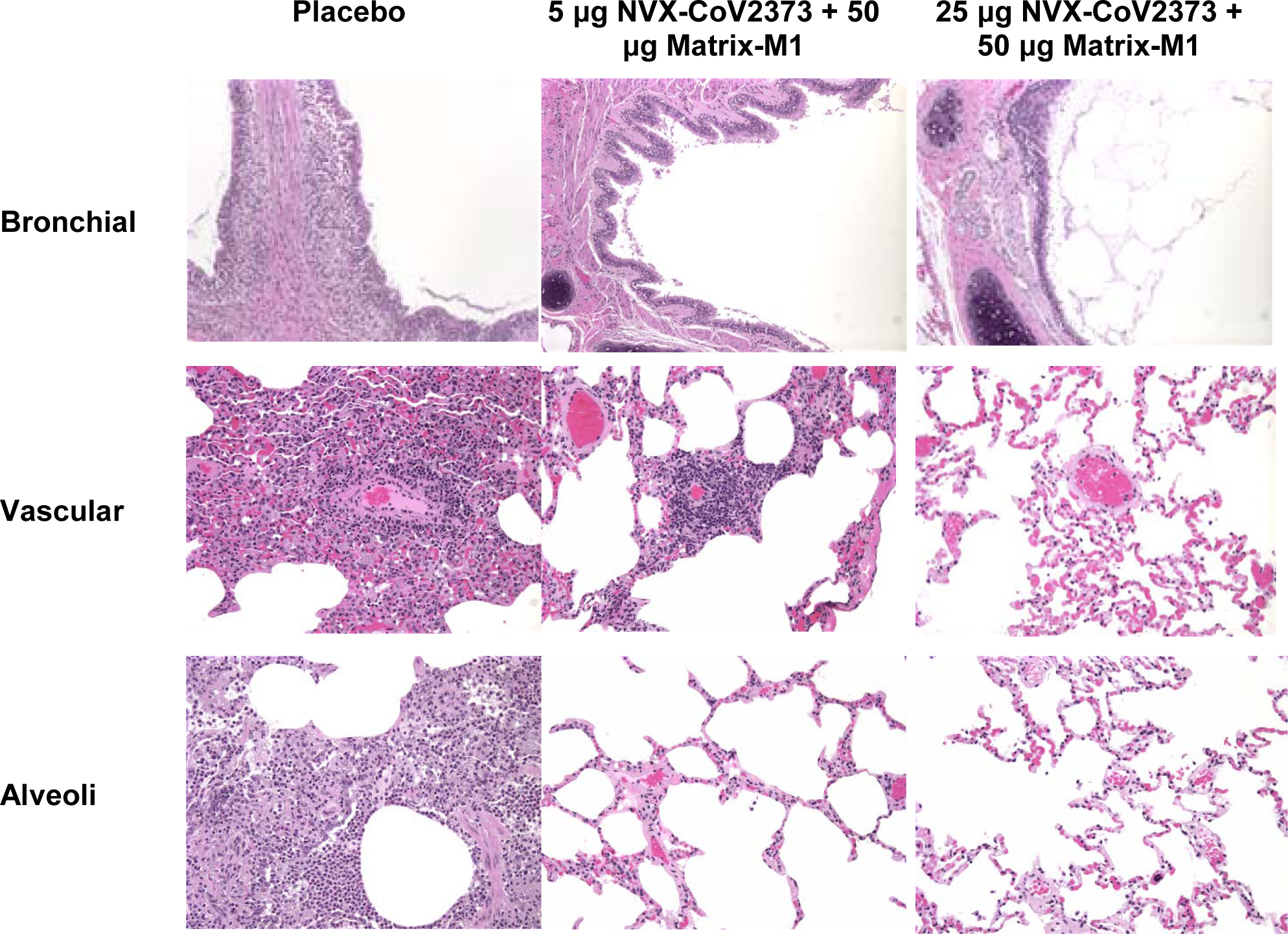
Representative histopathology of lungs from NVX-CoV2373 vaccinated cynomolgus macaques challenged with SARS-CoV-2 (WA1 strain). Histological findings of representative placebo treated animals included eosinophils expanding the mucosa of the bronchi, perivascular mononuclear infiltrate, with mixed macrophages and neutrophils within the alveoli at 7 days post infection. In the 5 μg dose group, one animal had mild to moderate perivascular infiltrate while other animals had no remarkable findings. There were no remarkable histological changes in the bronchial, vascular or alveoli in animals vaccinated with 50 μg NVX-CoV2373. There was no evidence of exacerbated lung inflammation in NVX-CoV2373 immunized animals.

## Discussion

Here, we report the immunogenicity and the protective efficacy of a prefusion, stabilized, full-length SARS-CoV-2 S vaccine (NVX-CoV2373) in the cynomolgus macaque model permissive to infection [11]. Prime and booster immunization with NVX-CoV2373 vaccine with Matrix1-M adjuvant induced high levels of anti-S IgG and antibodies that blocked SARS-CoV-2 spike protein binding to the hACE2 receptor and neutralized the virus. Importantly, vaccinated nonhuman primates had little or no detectable replicating virus (sgRNA) in either upper or lower respiratory tracks. These results demonstrate a potential of NVX-CoV2373 vaccine to protect the lower respiratory track against pulmonary disease and upper respiratory track against virus replication thus helping to establish herd immunity and to halt the COVID-19 pandemic and its devastating global impact.

## Acknowledgments

Funding for certain studies was provided by the Coalition for Epidemic Preparedness Innovations (CEPI), PO Box 123, Torshov, 0412 Oslo, Norway.

## Author contributions

MGX, GS, GG, JHT, ADP, MJM, MBF and LE contributed to conceptualization of experiments, generation of data and analysis, and interpretation of the results. NP, JHT, BZ, and SM performed experiments. MGX, NP and KL coordinated projects. GS, GG, MBF, PAP, MGX, NP and LE contributed to drafting and making critical revisions with the help of others.

